# Normative Modelling of Molecular-based Functional Neurocircuits Captures Clinical Heterogeneity Transdiagnostically in Neuropsychiatric Patients

**DOI:** 10.1101/2023.10.21.563428

**Authors:** Timothy Lawn, Alessio Giacomel, Daniel Martins, Mattia Veronese, Matthew Howard, Federico E. Turkheimer, Ottavia Dipasquale

## Abstract

Clinical neuroscience principally aims to delineate the neurobiology underpinning the symptoms of various disorders, with the ultimate goal of developing mechanistically informed treatments for these conditions. This has been hindered by the complex hierarchical organisation of the brain and extreme heterogeneity of neuropsychiatric disorders. However, recent advances in multimodal analytic techniques – such as Receptor Enriched Analysis of Connectivity by Targets (REACT) – have allowed to integrate the functional dynamics seen in fMRI with the brain’s receptor landscape, providing novel trans-hierarchical insights. Similarly, normative modelling of brain features has allowed translational neuroscience to move beyond group average differences between patients and controls and characterise deviations from health at an individual level. Here, we bring these novel methods together for the first time in order to address these two longstanding translational barriers in clinical neuroscience. REACT was used create functional networks enriched with the main modulatory (noradrenaline, dopamine, serotonin, acetylcholine), inhibitory (GABA), and excitatory (glutamate) neurotransmitter systems in a large group of healthy participants [N=607]. Next, we generated normative models of these networks across the spectrum of healthy ageing and demonstrated that these capture deviations within and across patients with Schizophrenia, Bipolar-disorder, and ADHD [N=119]. Our results align with prior accounts of excitatory-inhibitory imbalance in schizophrenia and bipolar disorder, with the former also related to deviations within the cholinergic system. Our transdiagnostic analyses also emphasised the substantial overlap in symptoms and deviations across these disorders. Altogether, this work provides impetus for the development of novel biomarkers that characterise both molecular- and systems-level dysfunction at the individual level, helping facilitate the transition towards mechanistically targeted treatments.

**Significance statement:** Human beings show enormous variability, with inter-individual differences spanning from neurotransmitters to networks. Understanding how these mechanisms interact across scales and produce heterogenous symptomatology within psychiatric disorders presents an enormous challenge. Here, we provide a novel analytic framework to overcome these barriers, combining molecular-enriched neuroimaging with normative modelling to examine neuropathology across scales at the individual level. Our results converge on prior neurobiological accounts of schizophrenia and bipolar disorder as well as the heterogeneity of ADHD. Moreover, we map symptomatology to molecular-enriched functional networks transdiagnostically across these disorders. By bridging the gap between dysfunctional brain networks and underlying neurotransmitter systems, these methods can facilitate the transition from one-size-fits-all approaches to personalized pharmacological interventions at the individual level.

## 1. Introduction

Neuropsychiatric disorders present a formidable healthcare challenge for which we remain largely bereft of meaningful treatments. Indeed, first line pharmacotherapies largely engage poorly understood pharmacological mechanisms discovered serendipitously decades ago (1) and remain ineffectual for many patients (2–4). The reasons for this are highly multifaceted, but ultimately reflect the extreme difficulty of understanding how complex aberrations of cognition and affect map onto underlying neurobiology. This in turn relates to the overarching challenge of understanding the brain, which is best described as a complex system whose diverse constituent components interact across scales and time to bring to bear the genetic and environmental interactions which collectively shape our experience and behaviour (5).

Neuropsychiatric disorders layer additional heterogenous pathophysiology on top of this inherent neurobiological complexity, resulting in significant variability in both neural mechanisms and symptomatology which show similarities across, and differences within, diagnostic boundaries (6–11). In other words, two individuals with a shared diagnosis may not have identical symptoms and two individuals with different diagnoses may share similar symptomatology. Furthermore, psychiatric conditions are highly co-morbid, adding to the challenge of disentangling underlying mechanisms (12–19). For this reason, many treatments are utilised trans-nosologically, with selective serotonin re-uptake inhibitors (SSRIs) having FDA approval for treatment of at least 10 different psychiatric disorders (20). In short, the current diagnostic paradigm is undermined by symptoms and treatments that are non-specific within and across disorders and that are poorly mapped onto underlying neurobiology (21). Despite this, the majority of research continues to largely employ a case-control-based paradigm, in which group average differences are characterised between clinical cohorts and matched healthy participants, inherently neglecting such heterogeneity. Precision psychiatry, and precision medicine more broadly, aims to move beyond large and poorly defined groups towards refined stratified or even individualised treatment based on underlying pathophysiological mechanisms (22). For example, the Research Domain Criteria (RDoC) framework, proposed by the National Institute of Mental health (NIMH), aims to re-examine mental disorders from the perspective of neurobehavioral functioning, regardless of conventional diagnostic categories (23–25). This is nicely exemplified by a null result for the primary outcome measure of a clinical trial. Currently, this reflects the average treatment response across all patients included in the study. However, the substantial heterogeneity within diagnostic groups may mean that in fact some patients do derive meaningful benefit (so called “responders”), whilst others show no real improvement (so called “non-responders”). Thus, the failure of the trial to demonstrate the clinical utility of a given treatment may be due to the inherent limitations of the intervention studied, but could also reflect an inability to effectively stratify, signifying the need for mechanistic biomarkers to enhance or supplant conventional diagnostic criteria within clinical practice. Whilst this theoretical line of argumentation is compelling, the practicalities of generating meaningful mechanistic biomarkers which can facilitate a transition to precision medicine has proven extremely challenging, with new tools required to characterise neuropathology at the subject level in a manner amenable to intervention.

Neuroimaging offers a non-invasive set of methods which can measure the structure and function of the brain, providing insights into the neurobiology underpinning psychiatric disorders. However, within conventional analytic frameworks, neuroimaging data is typically analysed for differences between patients and controls as well as for relationships between patients’ brain features and clinical measures. The latter offers insights into how variability across patients’ brains relates to severity of symptomatology. However, the ability to utilise these relationships across subjects to target treatment as well as transfer of these relationships to apply within additional patients from outside the original cohort remains limited. A set of emerging methods have been developed to address this specific issue. Normative modelling aims to robustly characterise what certain aspects of brain structure or function “should” look like across the spectrum of healthy ageing based on a set of predictor variables, typically demographics such as age and sex (26–30). This has been analogised as “growth charts for the brain” as it follows the same logic as predicting what a child’s height should be given their age and sex, except here the predicted measure is a particular facet of brain structure or function derived from neuroimaging. Once a model has been trained to make predictions about a given brain feature, it can be utilised to examine whether an individual with a given diagnosis, or particular constellation of symptoms, shows significant deviations from that expected given their demographics. To date, applications of normative modelling have largely been limited to structural imaging data, primarily due to its availability, practical simplicity, and interpretability. Normative models of structural grey and/or white matter measures have been explored in schizophrenia (31–34), bipolar disorder (32, 34), Alzheimer’s disease (35, 36), ADHD (32, 37), autism spectrum disorder (32, 38, 39), depression (32), and obsessive-compulsive disorder (32). A key emerging theme of this work to date is that whilst patients tend to have more extreme deviations than controls, the spatial distribution of these is extremely heterogeneous, to the extent that group average patterns of neuropathology are simply not representative of most individual patients. However, the exclusive application of normative modelling to structural imaging may also have constrained the potential insight that can be gained. Recent work has shown that these heterogenous structural deviations are often embedded within shared functional networks (32), emphasising the need to utilise functional imaging measures. Thus, despite the additional complexity, noise, and challenging interpretability, the rich spatiotemporal dynamics of functional imaging may offer benefits over the subtle and heterogeneous structural changes seen in psychiatric disorders.

Although structural imaging has been the focus of normative modelling thus far, and functional imaging may offer additional insights, successes in mechanistically stratifying patients based on functional neural measures remain scarce and no functional neuroimaging-based tool has been meaningfully exploited in clinical practice. One key reason underlying this is that fMRI offers only an indirect measure of neuronal function, indexing blood flow changes in voxels encompassing in the order of hundreds of thousands of neurones. As such, it remains abstracted from the cellular and molecular mechanisms that ultimately constitute brain function, and crucially, upon which interventions act. We contend that this may mean that deviations from health characterised using fMRI, as well as the aforementioned structural measures, are not readily amenable to targeted clinical intervention. However, a novel suite of analytic approaches which aim to incorporate micro-scale molecular information into the analysis of macro-scale fMRI dynamics offer critical opportunities to bridge the gap between these scales, providing new trans-hierarchical insights into brain function and dysfunction (see (40) for extensive review). Whilst much work has focussed on how bridging the gap between molecular and systems level mechanisms can help us better understand the inherent architecture of the brain, this work is especially well suited to provide novel biomarkers that link neuropathology through to pharmacotherapy in a mechanistic and data driven manner (40). For example, Receptor-Enriched Analysis of functional Connectivity by Targets (REACT) has proven useful in characterising the complex psychopharmacological effects of various drugs (40–43). Furthermore, it is increasingly being applied to clinical conditions, with recent proof of concept papers demonstrating its potential to stratify patients with osteoarthritis who may respond preferentially to placebo or duloxetine (Martins *et al.*, 2022) as well as normalise aberrant serotonergic processing associated with an emotional task in individuals with autism spectrum disorder (Wong *et al.*, 2022). This constitutes a substantial step towards mechanistic biomarkers that may offer meaningful capacity to link pathology to treatment.

Both normative modelling and molecular-enriched analyses offer substantial promise to overcome two of the pre-eminent limitations in biomarker development; complexity in the form of heterogeneity and the hierarchical organisation of the brain. As such, their combination offers a potential path forward for both mechanistic elucidation as well as to help bring neuroimaging closer to clinical implementation. To this end, here we utilised REACT to derive networks enriched within the main modulatory (noradrenaline, dopamine, serotonin, and acetylcholine), excitatory (glutamate), and inhibitory (GABA) neurotransmitter systems within two datasets. We subsequently generated normative models of these molecular-enriched networks across the healthy ageing spectrum. We then examined the clinical heterogeneity within and across patients suffering from schizophrenia (SCHZ), bipolar disorder (BPD), and attention-deficit hyperactivity disorder (ADHD). Only limited between group differences were observed using conventional diagnostic labels. Furthermore, between-subject similarity analyses revealed that deviation scores showed similarities within, but also across diagnostic groups. Moreover, a transdiagnostic deviation-symptom mapping approach yielded relationships between the cholinergic and glutamatergic systems to a dimensionally-reduced measure of symptomatology across all patient groups. Altogether, this offers significant progress towards generating novel biomarkers transcending organisational hierarchies of the brain and conventional diagnostic compartmentalisation, which in the longer term will offer a tantalising opportunity to link targeted treatment through to specific domains of neurobehavioral dysfunction.

## 2. Methods

### 2.1 Datasets

This study utilises two separate existing datasets. Firstly, the healthy ageing CamCAN dataset (obtained from the CamCAN repository available at http://www.mrc-cbu.cam.ac.uk/datasets/CamCAN/ (44, 45)) was chosen as it spans the full spectrum of healthy ageing. This allows normative models to provide relatively robust estimates across different test datasets, permitting some level of generalisability for subsequent investigations. Secondly, the UCLA phenomics dataset ((46), available from https://openneuro.org/datasets/ds000030/versions/1.0.0), was selected as it included healthy individuals as well as multiple psychiatric cohorts with deep clinical phenotyping, allowing for examination of deviations from healthy individuals both within and across conventional diagnostic boundaries. As both datasets contained extensive imaging, cognitive, and clinical testing, here we only describe measures utilised within the present study, with readers directed to the above resources for a comprehensive outline of these datasets.

### 2.2 Participants

#### CamCAN

The data used here is from the stage two of CamCAN, which contains a smaller subset of 700 individuals who met eligibility criteria to proceed from stage one, including 100 individuals from each decile (18-87 years old). These criteria included having no signs of diminished cognitive health (mini mental stage (MMSE) < 24, severe memory defects, or consent difficulties), no signs of communication difficulties (hearing problems, insufficient English language capabilities, or visual acuity difficulties), no self-reported medical problems (Parkinson’s disease, motor neurone disease, multiple sclerosis, cancer in the last 6 months, strike, encephalitis, meningitis, epilepsy, head injury, recent or uncontrolled high blood pressure, pregnancy, BPD, SCHZ, or psychosis), no mobility problems (restrictions that would prevent participation or inability to walk 10 meters), no substance abuse (past or current treatment for drug abuse, current drug usage, or refusal to answer questions regarding drug abuse), and no MRI safety contraindications (such as implanted devices, claustrophobia, or inability to lie still for an hour).

#### UCLA

All UCLA participants were aged between 21-50, had completed at least 8 years of formal education, had no significant medical illness, were adequately cooperative, had visual acuity 20/60 or better, tested negative for drugs of abuse (Cocaine; Methamphetamine; Morphine; THC; and Benzodiazepines), were not pregnant, were not left handed, did not have a history of head injury with loss of consciousness or cognitive sequelae, and did not have MRI contraindications (e.g. claustrophobia or metal in body). Participants in the healthy group [N = 130] were excluded if they had lifetime diagnoses of a psychiatric disorder. This included screening for sub-threshold ADHD using the Adult ADHD Interview and defined as 4 or more ADHD inattentive or hyperactive/impulsive symptoms in either childhood or adulthood. Each of the patient groups (SCHZ [N = 50], BPD [N = 49], and ADHD [N = 43]) excluded anyone with one of these other diagnoses. Stable medications were permitted for the patients.

Diagnoses for the clinical cohorts followed the Diagnostic and Statistical Manual of Mental Disorders (Fourth Edition)(47), utilising the Structured Clinical Interview for DSM-IV (SCID-I (48)) supplemented by the Adult ADHD Interview (a structured interview form derived from the Kiddie Schedule for Affective Disorders and Schizophrenia, Present and Lifetime Version (KSADS-PL)(49)). Interviewers were required to meet minimum standards of acceptable symptom agreement (overall kappa of .75, a kappa specificity of .75, kappa sensitivity of .75, and .85 kappa for diagnostic accuracy). Diagnostic and Symptom elicitation skill was also assessed with the SCID Checklist of Interviewer Behaviours (50) and the Symptom Checklist of Interviewer Behaviours (51). Ongoing quality assurance checks were conducted to ensure sufficient symptom agreement was being met.

### 2.3 Clinical and behavioural data

#### UCLA

A comprehensive list of the behavioural assessments can be found in table 3 of the original manuscript (46). Here, we utilised symptom measures from the Young Mania Rating Scale-C (YMRS), Hamilton Psychiatric Rating Scale for Depression (HAMD-17), Brief Psychiatric Rating Scale (BPRS), Hopkins Symptom Checklist (HSCL), and Adult Self-Report Scale v1.1

Screener (ASRS). Additional trait measures included in our analyses were Barratt Impulsiveness Scale (BIS), Scale for Traits that Increase Risk for Bipolar II Disorder (BPT), and the Chapman Scale for Perceptual Aberrations (CHAP). The acronyms here correspond to those used in figures throughout the manuscript. Symptom and trait scores selected had data available across all three clinical groups, providing measures of symptoms that are conventionally associated more with one of the diagnostic groups, but with potential involvement within each. Participants with incomplete demographic or psychometric data were excluded. Where relevant sub-scores were available for these symptom and trait measures, we utilised these within subsequent analyses to preserve the rich dimensionality of this phenotypic data. Where sub-scores were not available, we used the total summary score. In total, this offered 28 measures.

We additionally examined within-and between-group similarity of these scores. First, we plotted density curves of each score split by clinical diagnosis to see how overlapping or non-overlapping they were. Next, we created a correlation matrix which examined how correlated each subject was to every other subject across all 28 clinical scores available. This essentially offers a metric of between-subject symptom similarity.

### 2.4 Dimensionality reduction of psychometry

We examined the level of collinearity between the large number of different highly colinear clinical measures available within the UCLA dataset associated with the deep phenotypic characterisation and plotted these as a correlation matrix. Given the high levels of collinearity observed, we utilised principal components analysis (PCA, implemented in Python using sklearn) to reduce down the 28 different psychometric sub-scores and summary scores into a smaller set of components explaining a substantial portion of the variance in these scores across subjects. The psychometric data was normalised (using sklearn.preprocessing.StandardScaler) prior to being entered into the PCA. Importantly, we pooled together the scores across the three clinical cohorts in order to examine this symptomatology trans-diagnostically with the aim to identifying constellations of related symptoms. Components were retained based on the eigenvalue one criterion.

### 2.5 Imaging acquisition

#### CamCAN

All MRI datasets were collected at a single site (the Medical Research Council (UK) Cognition and Brain Sciences Unit) in Cambridge, UK) using a 3 T Siemens TIM Trio scanner (version syngo MR B17) with a 32-channel head coil. Functional MRI data were acquired with a T2* weighted GE EPI sequence, N=261 volumes of 32 axial slices 3.7mm thick, 0.74mm gap, TR=1970ms; TE=30ms, FA=78 deg; FOV =192 mm × 192 mm;3 × 3 x 4.44 mm, TA=8mins 40s. Additionally, a field map was acquired: PE-GRE, TR=400ms, TE=5.19ms/7.65ms, 1 Magnitude and 1 Phase volume, 32 slices 3.7mm thick, 0.74mm gap, FA=60deg, FOV= 192 × 192mm, 3 × 3 × 4.44, TA=53s. A 3D MPRAGE was also collected: TR=2250ms, TE=2.99ms, TI=900ms; FA=9 deg; FOV=256x240x192mm; 1mm isotropic; GRAPPA=2; TA=4mins 32s.

#### UCLA

MRI data were acquired on a 3T Siemens Trio scanner, located at the Ahmanson-Lovelace Brain Mapping Center (Siemens version syngo MR B15) and the Staglin Center for Cognitive Neuroscience (Siemens version syngo MR B17) at UCLA. Functional MRI data were collected using a T2*-weighted echoplanar imaging (EPI) sequence: slice thickness=4 mm, 34 slices, TR=2 s, TE=30 ms, flip angle=90°, matrix 64×64, FOV=192 mm, oblique slice orientation. An additional MPRAGE T1 was also collected: TR=1.9 s, TE=2.26 ms, FOV=250 mm, matrix=256×256, sagittal plane, slice thickness=1 mm, 176 slices.

### 2.6 Imaging pre-processing

The two datasets (CamCAN and UCLA) were used from their raw format and preprocessed for this work with nearly identical pipelines. FMRIPrep was used to undertake initial processing steps with subsequent denoising, filtering, and registration implemented separately (52). The only difference was the utilisation of a field map to correct for field inhomogeneity in the CamCAN data which was not performed for the UCLA data.

The fMRIPrep pipeline first corrected the structural T1 images for intensity non-uniformity and extracts the brain (i.e., skull stripping). This extracted brain was then spatially normalised into standard space through non-linear registration to the MNI152NLin6Asym template using ANTs (53). fMRI data were corrected for motion using MCFLIRT ((54)) and co-registered to the T1 weighted image using boundary-based registration (BBR) with six degrees of freedom (55). For CamCAN participants only, a subject-specific fieldmap was used to correct distortion caused by field inhomogeneity (56, 57). fMRI data were also slice-time corrected, smoothed with an isotropic Gaussian kernel of 6 mm FWHM (full-width half-maximum), and automatic removal of motion artifacts using independent component analysis (ICA-AROMA) was performed (58). Finally, using separate in-house scripts, subject-specific white matter (WM) and cerebrospinal fluid (CSF) masks were generated from segmentation of structural images, eroded to reduce partial volume effects with grey matter (GM), co-registered to the subject-specific functional space, and used to extract and regress out mean WM and CSF signals from each participant’s functional images. Finally, data were high-pass temporal filtered with a cut off frequency of 0.001 Hz and normalised to the standard MNI152NLin6Asym template space. Participants with head motion exceeding 0.5 mm framewise displacement were excluded (59, 60). All fMRIprep quality control images and metrics were visually inspected for every subject, with participants additionally excluded for significant dropout in fronto-temporal regions or repeated registration failures.

### 2.7 Population-based molecular templates

We employed transporter and receptor density maps from the noradrenergic, dopaminergic, serotonergic, cholinergic, and glutamatergic, and GABAergic systems. These are group average templates derived from healthy cohorts separate to the functional imaging datasets examined here. These have been widely utilised in our previous work (41–43, 61–65), as well as by the broader imaging community (66, 67). Here, we chose to use transporters for the neuromodulatory systems as these provide a general measure of the innervation and influence of a given receptor system over a given region (40). The NAT template was derived from 10 healthy participants utilising S, S-[11C]O-methylreboxetine (68). The DAT template was derived using 123I-Ioflupane single-photon emission computerized tomography (SPECT) images from 30 healthy subjects (HS) with no evidence of nigrostriatal degeneration (69). The SERT template was derived using [11C]DASB PET within 16 healthy individuals (internal PET database). The VAChT template utilised 18F-fluoroethoxybenzovesamicol PET within 12 healthy participants (70). [11C]flumazenil PET within 6 healthy individuals ((Myers et al., 2012) was used to produce the GABA-A template as described in (Dukart et al., 2018)). Finally, the mGluR5 template is from 31 healthy individuals using [(11)C]ABP688 (71). For each template, voxels within regions used as a reference for quantification of the molecular data in the kinetic model were replaced with the minimum value across all GM voxels in order to minimise the contribution of those regions without entirely excluding them from the models (i.e. occipital cortex for NAT and DAT as well as cerebellum for SERT, VAChT, and MGluR5). Finally, all templates were normalised by scaling image values between 0 and 1. To examine collinearity between the receptor systems, we calculated the correlation coefficients between each pair of PET templates as well as their Variance Inflation Factors (VIF). VIF quantifies the severity of multicollinearity in a least squares regression analysis 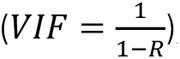, with higher values (i.e., above 5) denoting strong collinearity.

### 2.8 Receptor-enriched Analysis of functional Connectivity by Targets (REACT)

For each subject, functional networks associated with each of the molecular systems (NAT, DAT, SERT, VAChT, mGLuR5, and GABA-A) were estimated utilising a two-step multiple linear regression framework implemented in the REACT toolbox (https://github.com/ottaviadipasquale/react-fmri (72)). For further detailed methodological overview and discussion of REACT, see (40). The first regression analysis utilises the molecular templates as spatial regressors to calculate the dominant BOLD fluctuations associated with each molecular system. At this stage, both the fMRI data and the design matrix (i.e., the molecular templates) were demeaned and masked using a binarized atlas derived by combining a grey matter mask, the functional data across subjects, and PET templates in order to restrict the analysis to only GM voxels for which both functional and receptor density information was available. The second multiple regression analysis was performed to estimate the subject-specific molecular-enriched functional networks using the resultant subject-specific time series as temporal regressors. A binarized mask created by combining a grey matter mask with the functional data across subjects was used to restrict this GLM to only GM voxels for which there is functional data across all subjects. Once more, the time series estimated in the data and design matrix (i.e., the time series estimated in the first step) were demeaned, with the latter also being normalised to unit standard deviation. These were then averaged across participants to allow for visualisation of each resultant molecular-enriched network.

### 2.9 Parcellation

In order to limit computational burden, we parcellated our voxel-wise molecular-enriched networks using a custom combination of different atlases. For the cortex we utilised the Schaefer 400 region parcellation (73). We additionally added 15 regions from the Harvard-Oxford subcortical parcellation (left/right thalamus, left/right caudate, left/right putamen, left/right pallidum, left/right hippocampus, left/right amygdala, left/right accumbens, and brainstem)(74–77). Finally, we also utilised 28 cerebellar grey matter regions (all vermal and hemispheric lobules) from the SUIT atlas (78–80). Molecular-enriched FC values contained within each ROI were averaged, resulting in a total of 443 molecular-enriched network ROI values for each subject and each molecular system.

### 2.10 Normative modelling

In essence, this approach is used to generate individual molecular-enriched FC deviation maps based on a model of what an individual’s molecular-enriched network “should” look like, given their demographics. Previous experimental work has described various normative modelling approaches (31, 34–38, 38). As in most prior work, we implemented our normative models using the PCNtoolkit (version 0.27)(29). We chose to utilise a hierarchical Bayesian regression (HBR) approach which can accommodate signal and noise variance in data from multiple sites by estimating different but connected mean and variance components through shared prior distributions across sites (32, 81). A set of HBR models (82) were trained on healthy participants data from both the UCLA and CamCAN datasets, utilising a 70/30 train test split (implemented with sklearn.model_selection.train_test_split to ensure each dataset is equally represented in both training and test data), resulting in CamCAN = 346/150 and UCLA = 78/33 train/test subdivisions. By including both the CamCAN and UCLA data, this allowed us to generate a relatively stable distribution of estimates across the healthy ageing lifespan. A separate HBR model was estimated for each ROI and each receptor system (i.e. a separate model is fit for each region of the brain within each of the different molecular-enriched networks derived from our REACT and parcellation pipeline), using age and sex of participants to build a predictive model of molecular-enriched FC values. Specifically, these model site as a random effect and age and sex as fixed effects, resulting in normative regional molecular-enriched FC variance and uncertainty. Model performance was evaluated by the amount of variance explained when applied to the test sample. The subsequent analyses utilised only ROIs with positive explained variance (EV), in order to exclude areas where the model failed to converge on a stable estimate. Predictions using these models were also made for the clinical cohorts, where age and sex were used to predict their molecular-enriched networks. This provides both point estimates as well as measures of predictive confidence, allowing us to statistically quantify deviations from the normative molecular-enriched networks with regional specificity. Specifically, we computed a deviation *Z* score for each brain ROI, which describes the difference between the predicted molecular-enriched FC and the true molecular-enriched FC, normalised by the prediction uncertainty. In all subsequent analyses, we refer to the subset of healthy UCLA test subjects as HC_UCLA_.

### 2.11 Analysis of deviation scores between diagnostic groups

In line with the conventional diagnostic boundaries, we first examined whether the deviation *Z* scores differed between our three clinical cohorts and the HC_UCLA_ group. This was implemented ROI-wise as a 1X4 ANOVA within FSL’s Permutation Analysis of Linear Models program (PALM (83)) 2000 permutations and FDR correction). Lower-level t contrasts were used to delineate which groups were driving each significant higher-level result. To explore the added value of normative modelling, we repeated these analyses on the original molecular-enriched FC values for each network as would be done in a conventional REACT analysis.

We also collapsed down the deviation scores across ROIs for each molecular system to provide a single mean deviation score for each subject across their whole brain. This was motivated by the need for simple measures in order for putative biomarkers to be clinically practicable. We therefore used this singular metric which describes whether a given molecular system for a given subject is globally shifted more towards positive or negative deviations as well as the magnitude of this shift. We then conducted a 1X4 ANOVA for each molecular system to see whether these summary metrics differed between our three clinical cohorts and the HC_UCLA_ subjects. This analysis was implemented in Python (SciPy stats and statsmodels) with Tukey correction for lower-level comparisons. To further investigate the diagnostic capabilities of these summary metrics, we took significant between group differences from HC_UCLA_ at the lower-level and entered these mean *Z* values into a binary logistic regression analysis. Receiver operating characteristic curves (ROC) were plotted to display the trade-off between sensitivity and specificity. Area under the curve (AUC) was used to quantify predictive performance. This was also implemented in Python using statsmodels and sklearn.

### 2.12 Transdiagnostic analysis of deviation scores

We considered a novel approach to quantify how similar patterns of deviations were within and between groups. We speculated that whilst the HC_UCLA_ individuals would show a random pattern of deviations from normality, deviations within the patient groups may converge on similar patterns relating to a common disease process(es). We also expected the groups of patients to be more similar to one another than HC_UCLA_, signifying transdiagnostic similarity. Correlation matrices were generated for each molecular system, with each showing the correlation of deviation *Z* scores across all brain regions between each pair of individuals from the UCLA dataset (HC_UCLA_, SCHZ, BPD, and ADHD). This provides an overview of similarity across individuals, including both how similar the pattern of deviations is to those within the same group (within-group similarity) as well as to those in a different group (between-group similarity). We also compared the distributions of the within-group correlation coefficients across datasets using two-sample Kolmogorov-Smirnov tests (implemented with scipy.stats). Additionally, in order to explain this similarity metric further, we collapsed the within-group correlation matrix into one value of mean within-group correlation per subject, indicating how similar each subject is, on average, to the group they belong to. We then correlated this summary measure with the summary metric of deviation, to see whether a linear relationship exists between within-group similarity and magnitude of deviation from normality.

We also explored whether the within-group deviation similarity was related to the clinical scores, speculating that those who were highly similar may better reflect that diagnostic categorisation and that this would be reflected in their constellation of symptoms. This was implemented in Python and Bonferroni corrected for multiple comparisons across components and molecular systems (p < 0.05 / 24).

### 2.13 Transdiagnostic deviation-symptom mapping

The relationships between the four symptom-based PCs and the molecular-enriched network deviations from normality, expressed through the *Z* scores, was examined through mass univariate regression analyses. The resulting correlation coefficients reflect the magnitude of the statistical relationship between clinical and deviation data within each ROI across the brain. This was implemented through non-parametric permutation testing (2000 permutations and FDR correction) using PALM. In order to examine to what extent the normative modelling approach provides additional benefit, we also ran the exact same analysis on the raw molecular-enriched FC values, as would be done within a conventional REACT analyses.

## 3. Results

### 3.1 Demographics

Following quality control exclusions, imaging data of a total of 607 healthy subjects (CamCAN: N = 496, Mean age (SD) = 52.3 (18.3), M/F = 255/241; UCLA: N = 111, Mean age (SD) = 31.2 (8.7), M/F = 60/51) were included in the analysis. From the UCLA dataset, a total of 119 patients was used in the final analysis, including patients suffering from SCHZ (N = 38, Mean age (SD) = 35.1 (8.8), M/F = 32/6), BPD (N = 44, Mean age (SD) = 34.6 (8.9), M/F = 24/22), and ADHD (N = 37, Mean age (SD) = 32.2 (10.1), M/F = 19/18).

### 3.2 Between-subject symptom similarity

To first explore the phenotypic data, we examined each of the 28 different symptom scores within and across the different clinical groups. The distribution of scores for each of the groups overlapped for most symptoms, with only a few such as BPRS positive symptoms showing clear divergence across the conventional diagnostic groups (figure 1A). When examining how similar each patient’s constellation of symptoms was to every other patient, strong within-group similarity was seen for SCHZ and ADHD, whilst this was less clear for BPD (figure 1B). There are also clear examples of patients with different diagnoses who have strongly correlated symptomatology. Interestingly, BPD patients visually present greater within-group heterogeneity, with the group appearing divided into two sub-categories, i.e., either more similar to SCHZ or to ADHD, potentially indicative of it being an intermediate phenotype.

**Figure 1:**
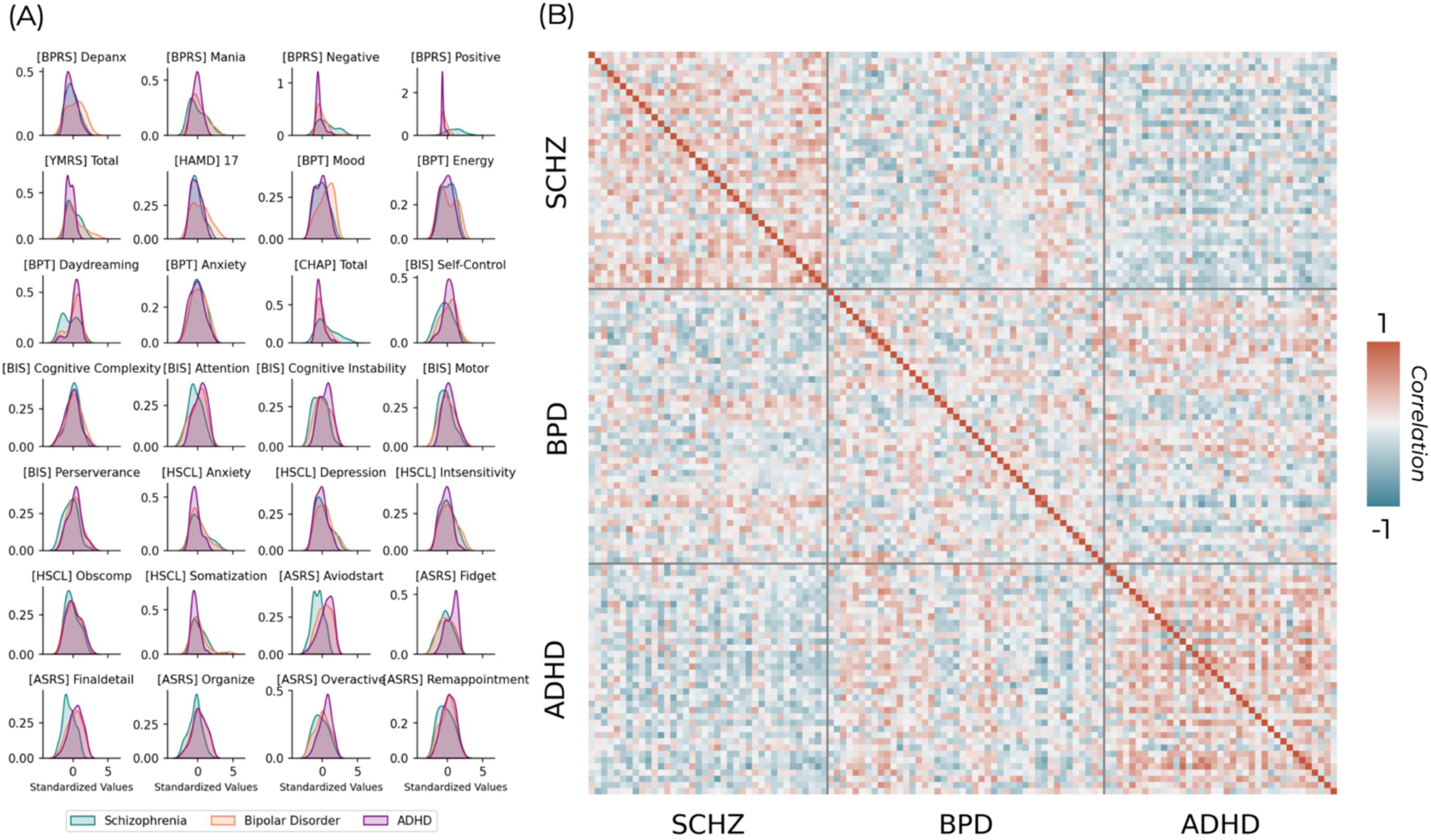
(A) Distributions of each psychometric measure for the different clinical cohorts. (B) Between-subject similarity matrix for these different normalised psychometric measures. Each position in the matrix represents the correlation coefficient between all psychometric scores for a pair of individuals. Grey bars separate the conventional diagnostic criteria, delineating matrix regions that show within (diagonal) and between group (off-diagonal) similarity.

### 3.3 Transdiagnostic psychometric dimensionality reduction

Across the 28 available psychometric scales and sub-scales there was generally strong overlap between clinical conditions. Cross correlations between these normalised psychometric measures showed a mixture of positive and negative relationships (figure 2A), with strongest values representing those which largely measure the same construct (e.g. BPRS depression and anxiety subscale correlates strongly with the total Hamilton depression score) and negative relationships between those measuring conceptually diverging constructs (e.g. the Adult Self-Report Scale primarily assesses ADHD symptomatology and shows mostly negative correlations with the Chapman Scales of psychosis proneness). The PCA run on these psychometric variables returned four variables which met eigenvalue one criterion and explained 68.9% of the total variance (figure 2B). These component scores retained significant overlap across the diagnostic groups (figure 2C), although PC2 showed a clearer separation across them than most of the original psychometric distributions (figure 1A).

**Figure 2:**
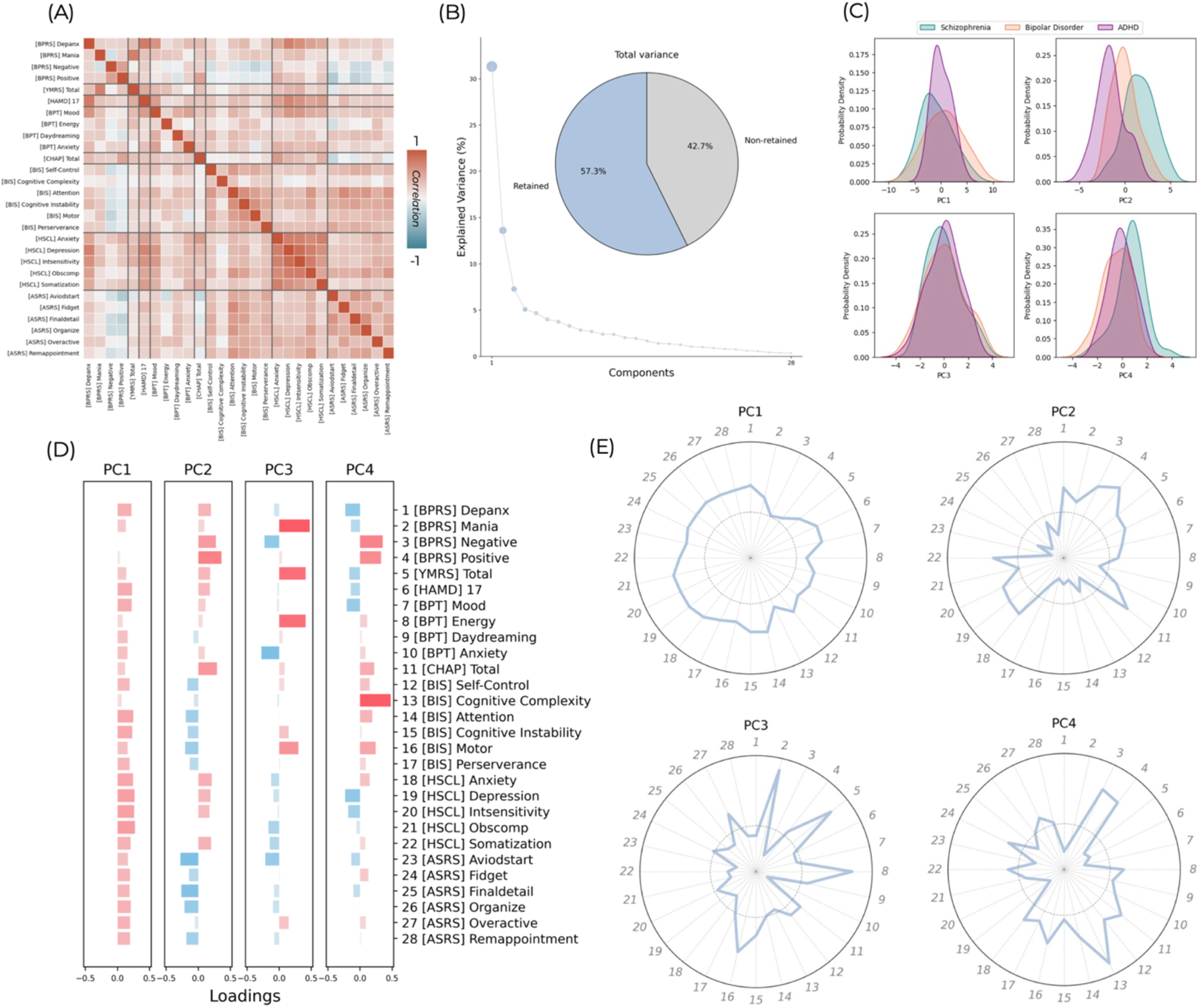
(A) cross-correlations between each pair of psychometric measures. (B) Component selection. Four components were retained which explain 68.9% of the variance in the original symptom scores. (C) The scores for each component across individuals within the three different clinical cohorts. (D) The loadings of the different symptom scores onto these four components shown as bar plots. These loadings are also shown as radar plots (E) for each component. The numbers 1-28 correspond to the symptom scores listed on the Y axis of plot D. The grey dashed inner circle represents a loading of zero, with portions of line outside this representing positive loadings and portions inside representing negative loadings.

The first principal component (PC1) represented general psychopathology with positive loadings across all psychometric measures (figure 2D/E). The subsequent 3 components showed more specific patterns of positive and negative loadings. PC2 seems to capture a generally psychotic and depressive symptomatology, with strongest loadings for positive symptoms, negative symptoms, psychosis proneness, as well as more moderate positive loadings for other measures such as for depression and anxiety. There were also moderate negative loadings across Barrett’s impulsiveness scale and the Adult Self-Report Scale. PC3 seems to be capturing a constellation of manic symptoms, with very strong loadings for mania (both the BPRS and YMRS), energy, and motor impulsiveness. Interestingly, there were also negative loadings for the Adult Self-Report Scale of ADHD symptoms, with the exception of overactivity. Finally, PC4 also captures psychotic symptoms, with strong loadings for positive symptoms, negative symptoms, and cognitive complexity but unlike PC2, the loadings for depressive symptoms are negative. Given the strongest loading is for cognitive complexity, this component may be separating patients that show relative preservation from cognitive decline. Additionally,, this component shows positive loadings across the impulsivity measures which are all negative in PC2.

### 3.4 Molecular-enriched networks

The REACT analysis, summarised in SI-figure 1A, produced a set of molecular-enriched networks for the CamCAN and UCLA healthy subjects as well as UCLA clinical cohorts. These networks are shown averaged across healthy individuals from both datasets in SI-figure 1B. The correlation coefficients between the different PET maps used in the REACT analysis (SI-figure 1C) as well as between the molecular-enriched networks (SI-figure 1E) were moderate, with generally stronger overlap within the neuromodulatory and excitatory/inhibitory neurotransmitter groups than between them. VIF values between the different PET maps were also moderate (SI-figure 1D), but importantly not surpassing the rule-of-thumb value of 5 above which issues of collinearity are considered sufficiently high to warrant additional consideration. Finally, the cross-correlations between the resulting molecular-enriched (SI-figure 1E) networks showed similar pattern to the cross-correlations to the PET maps from which they were derived (SI-figure 1C), again showing generally stronger overlap within the neuromodulatory and excitatory/inhibitory neurotransmitter groups than between them.

### 3.5 Modelling the normative molecular-enriched brain

Normative models were trained on 70% of the healthy control data across both the UCLA and CamCAN datasets and tested on the remaining 30%. These trained models explained a moderate amount of variance in molecular-enriched networks in this unseen data using age and sex as predictor variables. This EV varied substantially between molecular systems and across ROIs (figure 3A/B). Spatial distributions of explained variance largely reflected the key nodes of each molecular-enriched network. Herein, all analyses utilise only regions that had positive EV for that receptor system (from the total of 443 regions, this was 319, 307, 296, 257, 418, and 421 for NAT, DAT, SERT, VAChT, mGluR5, and GABA-A respectively). This reflects the spatial heterogeneity of the molecular-enriched networks, with each system being associated with key nodes in the brain and poorly associated with other regions, for which the normative models failed to converge on a stable estimate.

**Figure 3:**
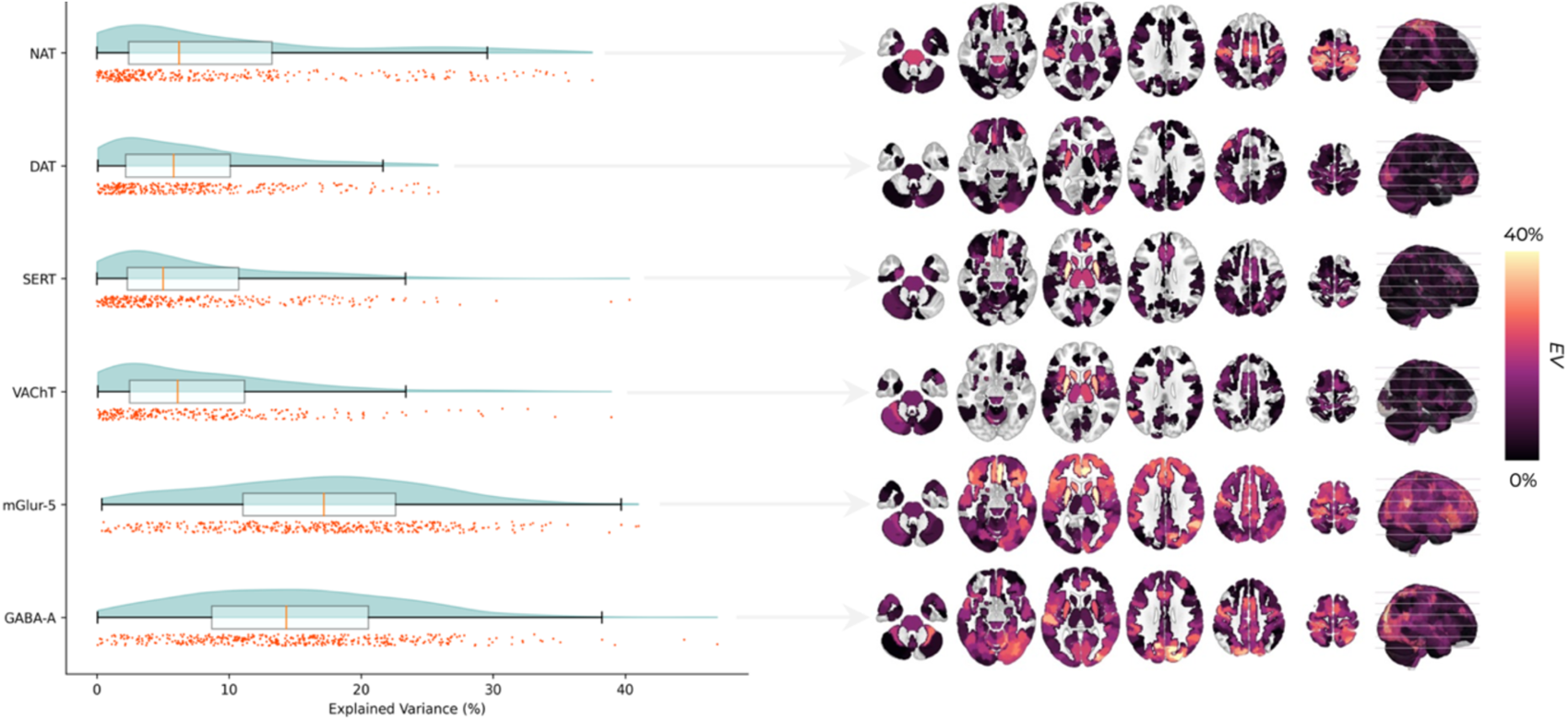
Models using age and sex as predictor variables explained a moderate amount of variance in the molecular-enriched networks for HC_test_ (left). Each raincloud represents the amount of variance explained across brain regions within each molecular-enriched system with each dot representing one brain ROI. The explained variance is also shown mapped back onto the brain for each molecular system (right).

### 3.6 Comparing deviations from normality across conventional diagnostic groups

The deviation scores (one *Z* value for every ROI, subject, and molecular system) were compared across groups for each molecular system. 1X4 ANOVAs revealed statistically significant differences within the VAChT and mGluR5 systems (figure 4B). Lower-level t-tests revealed that the higher-level VAChT result was driven by differences between HC_UCLA_ and SCHZ, whilst mGluR5 showed differences between HC_UCLA_ and both SCHZ and BPD (figure 4C). The VAChT system showed a mixture of regions where patients had more positive and more negative deviations than healthy controls, including the left putamen, left angular gyrus, bilateral precuneus, left supplementary motor area (SMA), right hemispheric lobule IX, and vermal lobule VIIIb. mGluR5 differences were largely in the direction of negative deviations within the SCHZ and BPD patients, with significant clusters in vermal lobule VI and right mid occipital cortex (figure 4C). It is also worth noting that the mGluR5 system had many clusters just below the significant threshold including the right insula (F = 6.13, p = 0.059 and SMA (F = 5.86, p = 0.059). No significant differences were found between HC_UCLA_ and the ADHD group. When considering differences between the clinical groups, lower-level comparisons additionally revealed that within the cholinergic system BPD patients had greater deviations than ADHD patients in the left precuneus and that SCHZ patients had greater deviations than ADHD patients within bilateral precuneus, left SMA, left angular gyrus, left pallidum, right hemispheric lobule IX, and vermal lobule VIIIb, similarly to what seen between SCHZ and HC_UCLA_. For the glutamatergic system, SCHZ patients showed a significantly greater deviation within the vermal lobule VI when compared to ADHD. Of note, we found no significant clusters within the conventional analysis (i.e. without normative modelling) running the same ANOVA on the molecular-enriched networks (SI-figure 2).

**Figure 4:**
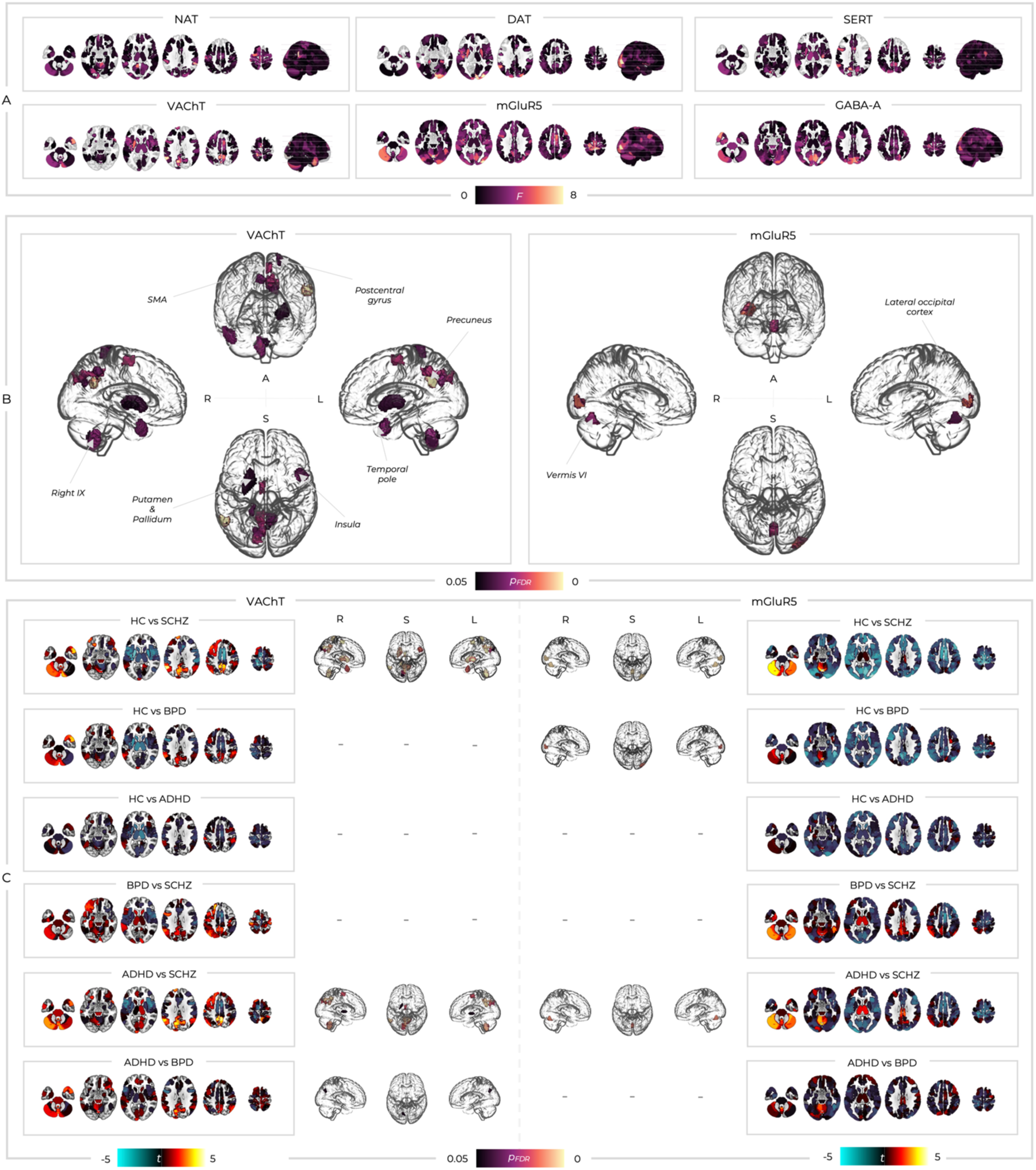
(A) The F statistics from a 1x4 ANOVA comparing analyses examining the deviation *Z* scores across the HC_UCLA_ and clinical groups (B) Only two molecular systems had significant (*p*_FDR_)between group difference (A; anterior view, L; left view, R; right view, S; superior view). SMA; supplementary motor area, STG; superior temporal gyrus. (C) Lower-level t-tests showing which between-group comparisons were driving the higher level results. t statistic colour bars correspond to the order of the sub-box titles such that when the left group has greater deviations these are blue and when the right group has greater deviations these are red/yellow (A; anterior view, L; left view, R; right view, S; superior view).

To examine whether we could produce a summary metric characterising each individual subject’s deviations within a given system rather than having one value for every ROI, we computed the mean *Z* values across regions and statistically compared these across the different groups of participants using 1x4 ANOVAs. We found significant differences across groups for the mGluR5 (F = 4.62, *p* < 0.001) and GABA-A (F = 3.24, *p* = 0.02) systems (SI-figure 3). Lower-level comparisons with Tukey correction revealed significant differences for mGluR5 only between SCHZ and healthy controls (*p =* 0.003) as well as BPD and healthy controls (*p* = 0.022). Similarly, for GABA-A lower-level comparisons showed significant differences for SCHZ compared to healthy controls (*p* = 0.014). The directionality of these differences was consistent with more negative average deviations within patients than controls, in line with the broadly negative t values across the brain seen in the ROI-wise analysis (see figure 4C). The mean *Z* values that showed significant differences in lower-level comparisons were then used within a binary logistic regression which revealed a moderate capacity to discriminate patients with SCHZ (SI-figure 3A/C) and BPD (SI-figure 3B) from healthy controls. Specifically, mean mGluR5 enriched-network deviation scores had an area under the curve (AUC) of 0.75 for SCHZ and 0.66 for BPD. Similarly, mean GABA-A enriched-network deviation scores resulted in an AUC of 0.73.

### 3.7 Transdiagnostic similarity of deviation scores

We then examined the between-subject correlations across the deviation scores from all ROIs for each molecular system to see whether they show similar within-and between-group patterns, as found in the analysis of the symptom scores reported in figure 1A. Between-subject correlations were calculated within each molecular system (figure 5A), providing a measure of how similar subjects’ deviations scores are across individuals in the same group and between different groups. The healthy controls were consistently dissimilar from each other, with the distribution of their between-subject correlation coefficients centred around zero (figure 5B). Kolmogorov-Smirnov tests revealed that the distribution of between-subject correlations in SCHZ and BPD were significantly different to those in HC_UCLA_ for all molecular systems (table 1), generally showing greater mean between-subject correlations in patients than HC_UCLA_ (SCHZ > BPD > ADHD > HC_UCLA_). The ADHD group showed significant differences from the HC_UCLA_ distribution for NAT and DAT, but not for the other molecular systems (table 1), although none of these ADHD results survived Bonferroni correction (p < 0.05 / 18). When considering relationships between these between-group similarity scores and summary mean deviation scores, we found significant negative correlations mGluR5 and GABA-A following Bonferroni correction (figure 5C). Participants showing greater within group similarity generally exhibiting more negative global deviations across their brain.

**Figure 5:**
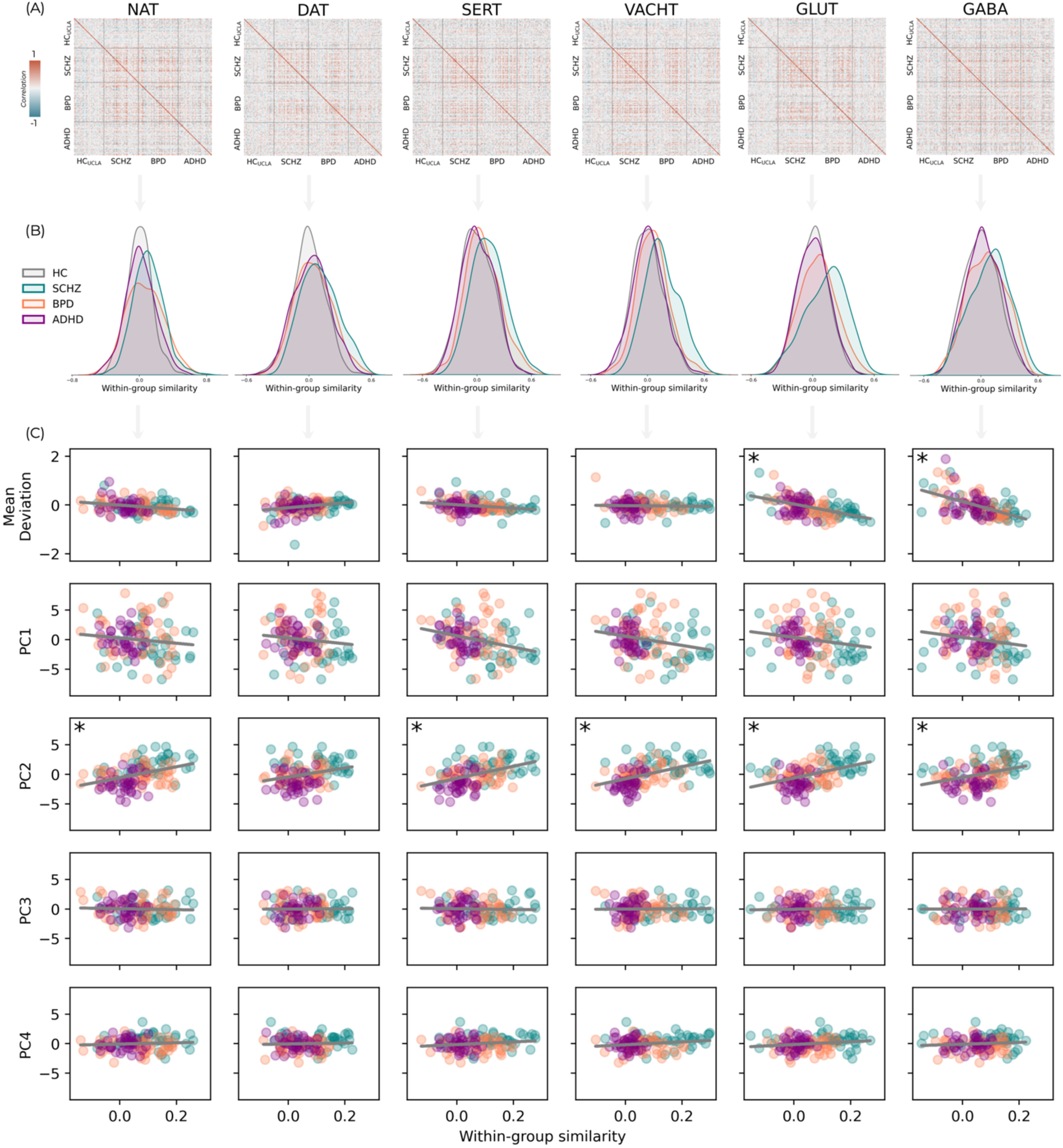
(A) Matrices of between-subject correlations of deviation *Z* scores across ROIs for each pair of individuals within and between groups. (B) The correlation coefficients for within group similarity displayed as density plots. (C) the relationship between patients similarity to the other patients in their diagnostic group and the overall deviation burden categorised by mean deviation *Z* score across their whole brain (top row) as well as each symptom component. Asterisks denote relationships that are significant following Bonferroni correction (*p* < 0.05/30).

**Table 1:** Within-group similarity for each receptor system and comparison across groups. Mean within-group similarity represents the mean correlation coefficient across every between-subject correlation within each group. Kolmogorov-Smirnov tests were conducted to compare these values from each clinical group to HC_UCLA._ Asterisks denote significant differences following Bonferroni correction (*p* < 0.05 / 18).

| Receptor | Group | Mean (SD) within-group similarity | KS statistic (vs HC) | p value (vs HC) |
| --- | --- | --- | --- | --- |
| NAT | *HC<sub>UCLA</sub>* | 0.00 (0.15) | - | - |
|  | *SCHZ* | 0.11 (0.18) | 0.27 | 0.00* |
|  | *BPD* | 0.06 (0.22) | 0.23 | 0.00* |
|  | *ADHD* | 0.02 (0.18) | 0.09 | 0.01 |
| DAT | *HC<sub>UCLA</sub>* | 0.00 (0.15) | - | - |
|  | *SCHZ* | 0.10 (0.18) | 0.24 | 0.00* |
|  | *BPD* | 0.05 (0.18) | 0.13 | 0.00* |
|  | *ADHD* | 0.03 (0.17) | 0.10 | 0.01 |
| SERT | *HC<sub>UCLA</sub>* | 0.00 (0.18) | - | - |
|  | *SCHZ* | 0.13 (0.19) | 0.26 | 0.00* |
|  | *BPD* | 0.07 (0.19) | 0.14 | 0.00* |
|  | *ADHD* | 0.01 (0.18) | 0.05 | 0.54 |
| VACHT | *HC<sub>UCLA</sub>* | 0.00 (0.16) | - | - |
|  | *SCHZ* | 0.15 (0.18) | 0.35 | 0.00* |
|  | *BPD* | 0.06 (0.18) | 0.15 | 0.00* |
|  | *ADHD* | 0.01 (0.17) | 0.05 | 0.53 |
| mGluR5 | *HC<sub>UCLA</sub>* | 0.00 (0.15) | - | - |
|  | *SCHZ* | 0.13 (0.20) | 0.35 | 0.00* |
|  | *BPD* | 0.05 (0.18) | 0.14 | 0.00* |
|  | *ADHD* | 0.01(0.15) | 0.03 | 0.90 |
| GABA-A | *HC<sub>UCLA</sub>* | 0.01 (0.18) | - | - |
|  | *SCHZ* | 0.10 (0.21) | 0.22 | 0.00* |
|  | *BPD* | 0.06 (0.20) | 0.14 | 0.00* |
|  | *ADHD* | 0.03 (0.18) | 0.05 | 0.38 |

We also examined whether these principal components related to the level of within-group similarity. Significant correlations following Bonferroni correction were found between PC2 and all receptor systems except for DAT, which were all consistent with greater similarity within group being related to greater PC2 scores (figure 5C and table 2). By visually inspecting the scatterplots, we can see that this was largely driven by the SCHZ group which generally shows greater within-group deviation similarity values and PC2 scores, whilst ADHD shows the inverse. Deviations in the SERT enriched network showed the opposite relationship with PC1, such that greater deviation similarity was associated with lower PC1 scores. Finally, we also examined the relationship between our summary mean *Z* values and the principal component scores. The only significant correlation was for the glutamatergic system and PC2 (r = -0.21, p = 0.024) which reflects our ROI-wise results, although this did not survive Bonferroni correction (p < 0.05 / 6).

**Table 2:** Correlations between within-group similarity and the four principal components for each different molecular system. Asterisks denote significant differences following Bonferroni correction (*p* < 0.05 / 30).

| Receptor | Component | Pearson's r | p value |
| --- | --- | --- | --- |
| NAT | PC1 | -0.13 | 0.170 |
|  | PC2 | 0.40 | 0.000* |
|  | PC3 | -0.05 | 0.606 |
|  | PC4 | 0.08 | 0.395 |
| DAT | PC1 | -0.12 | 0.189 |
|  | PC2 | 0.28 | 0.002 |
|  | PC3 | 0.01 | 0.900 |
|  | PC4 | 0.05 | 0.595 |
| SERT | PC1 | -0.28 | 0.002 |
|  | PC2 | 0.46 | 0.000* |
|  | PC3 | -0.04 | 0.679 |
|  | PC4 | 0.16 | 0.077 |
| VACHT | PC1 | -0.24 | 0.009 |
|  | PC2 | 0.47 | 0.000* |
|  | PC3 | 0.02 | 0.853 |
|  | PC4 | 0.18 | 0.045 |
| mGluR5 | PC1 | -0.18 | 0.044 |
|  | PC2 | 0.45 | 0.000* |
|  | PC3 | 0.04 | 0.652 |
|  | PC4 | 0.17 | 0.060 |
| GABA-A | PC1 | -0.17 | -0.17 |
|  | PC2 | 0.35 | 0.000* |
|  | PC3 | 0.01 | 0.932 |
|  | PC4 | 0.10 | 0.0283 |

### 3.8 Deviation-symptom mapping

We then examined the relationships between the four transdiagnostic symptom PCs and *Z* scores within each ROI and molecular system. Varying patterns of both positive and negative deviation-symptom relationships were seen across molecular systems and components (figure 6A). Across components and systems, only the relationships between PC2 and the VAChT and mGluR5 systems were significant (figure 6B). VAChT showed significant relationships with PC2 spanning bilateral insular and opercular regions as well as the left mid-cingulate, SMA, and precuneus. The mGluR5 systems showed relationships with PC2 spanning similar bilateral insular and opercular regions, as well as the bilateral superior temporal gyri (STG), bilateral cuneus, left lingual gyrus, bilateral mid-cingulate, and bilateral SMA. Both of these relationships were negative, such that higher PC2 symptom scores were associated with more negative *Z* scores (figure 6C). Additionally, the different subgroups of patients showed some degree of separation within these relationships, with SCHZ patients tending to have higher symptom scores and more negative deviations across both VAChT and mGluR5 systems. We also repeated this analysis using the original molecular-enriched networks to examine whether the normative modelling approach offered additional benefits. Only the VAChT system showed positive results, with similar but more limited significant clusters. Specifically, these were in the right insular and left SMA as described above, but also bilateral PFC. No significant results were found for mGluR5 in this basic analysis.

**Figure 6:**
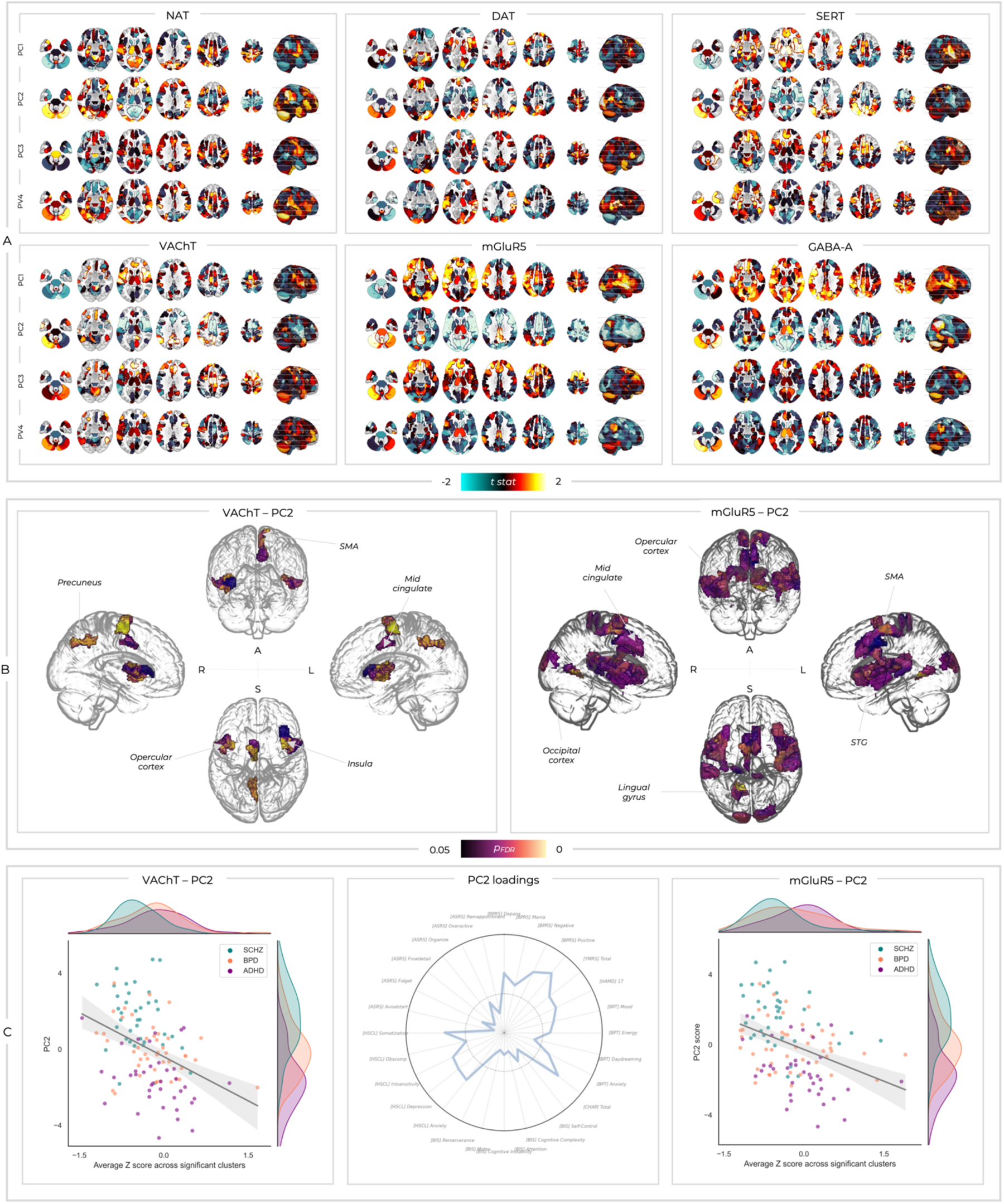
(A) The t statistics from multiple-linear regression analyses examining the relationship between deviation *Z* scores and the four principal components at each ROI. (B) Only two of these deviation-symptom relationships were significant. Significant *p*_FDR_ values are shown for the relationships between PC2 and deviations within the VAChT-and mGluR5-enriched networks (A; anterior view, L; left view, R; right view, S; superior view). SMA; supplementary motor area, STG; superior temporal gyrus. (C) The same relationships shown in the brain plots, with Z values averaged across the significant clusters and correlated with PC2 symptom scores. The loadings for PC2 are shown again here for context.

## 4. Discussion

Here, we bring together two novel classes of neuroimaging analytics which hold significant potential to circumvent key barriers to the neurobiological characterisation, and mechanistically informed treatment, of neuropsychiatric disorders. Our normative models performed comparably to previous work on structural neuroimaging data, but additionally captured both functional and molecular facets of the brain. Normative modelling revealed more robust between-group differences and deviation-symptom relationships than conventional REACT analyses, emphasising the power of describing neuropathology as divergence from estimates of healthy ageing. Our between-group findings converge to broadly implicate the excitatory-inhibitory balance in SCHZ and BPD, the cholinergic system in SCHZ, and largely failed to find specific disorder-based alterations in ADHD. However, moving beyond these categorisations, we found substantial similarity between individuals transdiagnostically and that deviations mapped onto symptom scores across groups. Interestingly, these transdiagnostic deviation-symptom relationships revealed separation of the three cohorts along the PC2 symptom scores as well as deviation scores within the glutamatergic and cholinergic systems. Herein, we discuss these key findings in light of the challenges presented by the hierarchical organisation of the brain and heterogeneity within clinical neuroscience.

### 4.1 Normative modelling of molecular-enriched functional networks

To our knowledge, this is the first time normative modelling has been directly applied to functional imaging data. Despite departing from the typical use of structural imaging features, our models explained a comparable amount of variance to those published previously. This constitutes a significant step forward for normative modelling, suggesting that this nascent field can begin to additionally exploit the rich spatiotemporal dynamics of functional imaging, which may offer benefits over-and-above measures of grey and white matter in characterising neurobiological heterogeneity. Indeed, the field of neural fingerprinting (84, 85) highlights the significant individual-specific content of resting-state fMRI data, which is being increasingly applied to describe clinical heterogeneity (86). Moreover, the number and size of openly available rs-fMRI datasets has increased markedly in the last decade, providing opportunity to explore different functional imaging metrics across a variety of clinical populations. Moving forward, side-by-side comparison of normative models utilising common structural approaches (such as voxel-based morphometry) alongside simple voxel-wise estimates of connectivity (such as global brain connectivity) may prove instructive in delineating the relative pros and cons of these different imaging modalities. Interestingly, deviations from these molecular-enriched networks revealed more robust results than conventional analysis on the molecular-enriched FC values, additionally highlighting the utility of characterising neuropathology and symptomatology as deviations from estimated normality.

Our results move beyond both conventional structural and functional MRI measures, which are inherently incapable of providing insights into their cellular and molecular underpinnings. The use of molecular-enriched functional imaging approaches offers a tantalising opportunity to circumvent these barriers and make use of already abundant fMRI data to provide insights spanning the molecular and systems levels (40). Crucially, this enables a non-invasive characterisation of dysfunction that is directly amenable to targeted pharmacotherapeutic intervention in a scalable manner. This could be achieved through a broad categorisation of which molecular-enriched networks are altered by acute drug challenges with different compounds, allowing these to be targeted in patients showing substantial deviations within networks enriched with the same molecular targets. Whilst we utilised molecular-enriched networks here, additional methods such as mapping patterns of deviation onto whole-brain transcriptomics may also prove powerful tools to this end. This work offers only a provisional proof-of-concept set of results, with more refined approaches deployed within more comprehensive datasets which include treatment responsiveness required to meaningfully test this idea.

### 4.2 Analyses using conventional diagnostic categories align with previous accounts of neuropathology

Our between-group analyses broadly converged upon the glutamatergic and GABAergic systems, potentially reflecting excitatory-inhibitory (E/I) imbalance, which is a widely implicated facet of neuropathology in SCHZ and BPD (87), and providing confidence in our methods. Across all our analyses comparing deviations between SCHZ and controls we identified differences in the glutamatergic and/or GABAergic systems, with patients broadly showing more negative deviations and greater similarity to one another than healthy individuals. Within SCHZ, various aspects of glutamatergic and GABAergic (dys)function are evidenced by key risk loci within genetic studies (88–91), altered resting gamma power (92–95), sensory gating deficits (96–98), changes in various TMS-EEG paradigms (99), reduced mismatch negativity amplitude (100, 101), reduced hippocampal of NMDA and GABA-A receptor density (102, 103), lower post-mortem levels of the GABA synthesising enzyme glutamate decarboxylase 67 (104–106), and parvalbumin positive interneurons in SCHZ (107–111). Although some of these measures have also shown null findings in additional studies, this critical mass of literature strongly implicates E/I imbalance within SCHZ. We further add to this overall picture with results relating to both glutamate and GABA, lending credence to the idea that E/I imbalance in SCHZ emerges from an interplay of these two systems, as opposed to one or the other, which remains an outstanding issue within the field (112). Similarly, the glutamatergic system was consistently implicated in BPD across our analyses. Despite a heavy focus on monoamines for over half a century, the glutamatergic system is increasingly thought to play a key role within mood disorders (113). A meta-analysis of magnetic resonance spectroscopy studies found that glutamate levels were elevated in BPD compared to controls when all brain areas were combined, regardless of medication status (114). A follow up meta-analysis largely corroborated this, identifying increased frontal glutamate and decreased mismatch-negativity (115). Additional evidence from genetics (116–124), blood and urine markers (125–128), as well as post-mortem studies (129–134) point towards glutamatergic dysfunction. We found no significant differences between glutamatergic or GABAergic deviations within SCHZ and BPD, whilst SCHZ and ADHD showed differences for both mGlur5 and VAChT which mirrored differences between SCHZ and controls. This suggests a general neurobiological similarity between SCHZ and BPD whilst ADHD participants more closely resembled controls. Our results are therefore broadly convergent with previous accounts of E/I imbalance in these disorders as well as some level of shared underlying neuropathology, offering confidence that our normative modelling approach aiming to link molecular-and systems-level mechanisms is capturing meaningful aspects of neuropathology. Furthermore, the failure of our conventional REACT analyses to identify any between group differences emphasises the value of describing pathology as divergence from normative values. Moreover, the fact that these differences were captured by whole-brain mean deviation summary scores provides tentative support that this approach could be used to create simplified clinical readouts of the integrity excitatory and inhibitory molecular-enriched networks.

The cholinergic findings relating to SCHZ throughout our results are intriguing. All currently approved antipsychotics target the dopamine D2 receptor (135), with actions on ventral tegmental area (VTA) dopaminergic neurons thought to underlie therapeutic efficacy in reducing positive symptoms whilst unwanted action on the substantia nigra results in extrapyramidal side effects (136–139). However, current treatments provide little benefit for negative or cognitive symptoms and many patients continue to experience residual positive symptoms or remain treatment resistant (140, 141). There is increasing interest in novel cholinergic compounds being able treat SCHZ symptoms, including positive, negative, and cognitive symptoms, whilst not being associated with the long-term side effects of dopaminergic antipsychotics (142, 143). Anticholinergic drugs can induce or exacerbate confusion, delirium, cognitive impairment, and hallucinations, with this mimicry of psychiatric symptoms indicative of the role cholinergic mechanisms may play in these symptoms when arising clinically (142, 144). Conversely, in the 1990’s xanolemine, a muscarinic antagonist being developed for cognitive symptoms of Alzheimer’s disease, was noted to have antipsychotic effects (145), and these results have been replicated in SCHZ patients (146–148). Interestingly, these studies showed benefits for positive and negative symptoms, but also cognitive performance. Crucially, xanolemine’s efficacy is not mediated by direct action on D2 receptors (149). Therefore, pharmacological manipulation of the cholinergic system has been robustly demonstrated to induce or ameliorate the symptoms of SCHZ. Our findings add credence to this view, suggesting that the cholinergic system is related to altered systems level dynamics in SCHZ compared to controls and that individuals with more negative cholinergic deviations had greater PC2 scores, which reflect core SCHZ symptoms. It is therefore tempting to speculate that xanolamine may act to normalise the cholinergic dysfunction spanning molecular and systems levels identified here. This may be mediated through indirect actions of the laterodorsal tegmentum on the dopaminergic system (150, 151), potentially preferentially acting on VTA dopaminergic neuron firing whilst sparing the substantia nigra-mitigating extrapyramidal side effects and tardive dyskinesia (152). However, additional preclinical evidence supports the idea that xanolemine may act through modulating glutamatergic microcircuits, which may in turn also impact dopaminergic transmission (153–155). These complex receptor sub-type-specific mechanisms have been discussed at length recently (142, 143, 156). However, a key point is the emerging idea that novel cholinergic compounds could provide antipsychotic effects through conventional dopaminergic pathways thought to drive positive symptoms, but additionally benefit for negative and cognitive symptoms not ameliorated by current treatments (142, 143). Our PC2 relates to a broad range of psychotic symptoms, potentially supporting links through to the cholinergic system regardless of diagnosis.

The lack of results for ADHD throughout our analyses is generally consistent with this disorder being extremely heterogeneous. Indeed, previous work utilising normative models of grey and white matter volume found that almost no brain regions showed consistent deviations within ADHD patients (37) and a recent meta-analysis of 96 structural and functional imaging studies in ADHD found a lack of regional convergence (157). Similarly, we identified no significant differences between ADHD subjects and healthy controls within any brain region nor in our summary measures of mean deviation scores. Intriguingly, we did find significant differences in the distribution of between-subject similarities for ADHD participants and controls for the noradrenergic and dopaminergic-enriched networks, although distributional differences were small and did not survive Bonferroni correction. The catecholamines are long implicated in the pathophysiology and treatment of ADHD (158, 159), adding credence to these similarity analyses which may offer an alternative lens through which to view the pattern of deviations across subjects. Analogous to the recent combination of normative modelling and network-lesion mapping, wherein deviations are examined for co-localisation to common networks (32, 160), it would be interesting to examine whether deviations within ADHD map onto regions strongly influenced by catecholaminergic transmission. This could be indexed through connectivity patterns of the noradrenergic locus coeruleus and dopaminergic ventral tegmental area, the distribution of different catecholaminergic receptors and transporters, or some multi-modal combination of these converging measures. A larger sample focussed explicitly on characterising ADHD deviations and symptomatology would be better placed to investigate this possibility.

### 4.3 Recharacterizing our analyses across diagnostic boundaries

Despite receiving different diagnoses, we found transdiagnostic similarities across groups within our analyses. Patients symptom scores were broadly correlated regardless of diagnosis, indicative of highly overlapping clinical phenotypes. This was substantially recapitulated in the deviation scores, with similarity matrices across all patients revealing strong correlations within the SCHZ and BPD groups as well as across them, especially within the glutamatergic and GABAergic systems. ADHD participants showed less clear within-and between-group similarity, more closely resembling the control group. These within-group similarity analyses may also relate to a general body of literature which suggests that neuropathology associated with various disorders of the brain may restrict the repertoire of network states a brain can inhabit (161–163). The level of within-group deviation similarity for glutamatergic and GABAergic systems was also inversely related to summary mean deviation scores, indicating that those who were more similar had a more negative mean deviation from normality across the brain. This was not as pronounced for the other systems, suggesting that the excitatory and inhibitory tone is perturbed more globally, whilst the cholinergic system, which was implicated in other analyses, may have more focal effects. Additionally, it is possible that cholinergic deviations are less consistent in directionality, with positive and negative deviations cancelling each other out within the summary mean *Z* measure. This idea is also supported by the fact that our mean glutamate *Z* score summary metric was related to PC2 transdiagnostically, although this did not survive Bonferroni correction, whilst no relationship was seen for the cholinergic system. Thus, these summary scores may prove particularly useful as biomarkers for excitatory and inhibitory neurotransmitter dysfunction. The between-subject similarity findings additionally underscore the challenges of clinical diagnostic categorisation, with clear within-group similarity but also significant correlation between clinical groups, reflecting the fact that current labels do seem to capture underlying neurobiology, but not with sufficient precision to fully differentiate groups or robustly target treatment. When considering symptomatology, we found relationships linking our dimensionally-reduced components through to deviations transdiagnostically. Our PC2 seemed to capture primarily aspects of psychosis, with strongest loading for positive symptoms, negative symptoms, and perceptual aberrations. Accordingly, our significant deviation-symptom mapping results revealed that the significant correlations between PC2 and cholinergic and glutamatergic network deviations also had delineation of the different clinical cohorts, with SCHZ patients generally showing higher PC2 scores and more negative deviation *Z* scores. The ADHD cohort displayed the converse pattern, generally exhibiting negative PC2 scores and deviation *Z* scores closer to zero. However, the fact that those ADHD patients with greater scores on the PC2 symptom axis also generally showed more negative deviations in the glutamatergic and cholinergic systems, as seen within SCHZ and BPD, highlights the utility of mapping symptoms to mechanisms transdiagnostically. Intriguingly, whilst BPD patients showed an intermediate distribution between SCHZ and ADHD in PC2, their deviation *Z* scores were more similar to SCHZ patients for the glutamatergic system and ADHD patients for the cholinergic system. This may reflect a molecular-functional distinction between groups, with the cholinergic system more related to SCHZ, whilst the glutamatergic system being implicated in both SCHZ and BPD. This offers an interesting way to examine similarities and differences between conventional diagnostic groups within this transdiagnostic symptom-network space. The fact that overall SCHZ and BPD appear to have more similar patterns of deviation across these molecular-enriched networks as well as occupy a more overlapping region in the PC2-glutamate/acetylcholine space to one another than to ADHD, but also have key differences between them, aligns with clinical presentation as well as aetiological theories of these disorders (87, 164, 165).

### 4.4 Neurotransmitter specificity and interactions

It remains unclear why we found cholinergic and not dopaminergic results across these analyses, given the core role dopamine is thought to play within the pathophysiology of SCHZ (136–139). The distribution of dopaminergic and cholinergic transporter density strongly overlaps within key regions such as the striatum. It is therefore entirely possible that our cholinergic results additionally reflect dopaminergic mechanisms, especially given how strongly mechanistically intertwined these systems are in shaping cortico-striato-thalamic circuitry (Lester et al., 2010). Moreover, there was additional high VAChT density within the thalamus, another region implicated in the pathophysiology of SCHZ (Pergola et al., 2015; Pinault, 2011), potentially contributing to the cholinergic findings. Regardless of the precise delineation between the two systems, these regions which differ between groups and relate to transdiagnostic symptomatology are influenced by the cholinergic and dopaminergic systems, with neurobiologically plausible accounts for cholinergic dysfunction giving rise to multiple symptoms in SCHZ. Future methodological development to more carefully tease apart relationships where receptor density and pathophysiology are highly colocalised will be crucial to truly link these measures through to treatment. Subsequent work could consider examining deviations within dopaminergic, cholinergic, glutamatergic, and GABAergic receptor subtypes to allow for clearer delineation between putative drug targets and core symptom domains. Alternatively, focussing on results which converge with other methodologies such as using seed-based connectivity from dopaminergic and cholinergic nuclei may be beneficial. Additionally, the availability of treatment response outcome measures would allow for the direct examination of whether dysfunction in a given molecular system is indeed predictive of treatment responsiveness to an intervention which targets that system. Methodological progress on this front will be crucial to move beyond simplified accounts of neurobiology and neuropathology, and consider the full complexity of multiple neurotransmitter systems acting in concert.

## 5. Limitations

This work is not without limitations. As with all molecular-enriched network analyses, we utilise group average PET templates acquired in separate healthy subjects. The validity of this approach has been described extensively within the broad literature utilising PET and transcriptomic data within neuroimaging, as discussed at length in (40). Several of these molecular systems also show moderate collinearity, which requires careful consideration in the multiple-regression approach employed within REACT. However, we show that VIF values are below 5, suggesting that levels are not broadly problematic in the models employed here. This does present a barrier to the inclusion of additional receptor systems in the future, and finding new ways to examine the full repertoire of molecular systems and receptor sub-systems within the human brain will be important in the longer term. Moreover, the cholinergic and dopaminergic PET templates utilised here share core regions of high transporter density, making the disentangling their contributions challenging. The cross-sectional nature of our data also confines us to observing associations rather than establishing causal relationships between deviations and clinical measures. Longitudinal studies are imperative to delineate the true temporal trajectory of molecular-enriched networks and their relationships to symptomatology as well as characterise their potential diagnostic, prognostic, and treatment predictive capabilities. Similarly, the sample sizes of the clinical cohorts utilised here were relatively small, with our proof-of-concept results requiring follow up in more robustly powered samples, ideally also including a diverse set of patients which often fall outside the scope of the strict selection criteria of smaller studies. Additionally, we only tested our deviation-symptom mapping in one dataset. Future work will require a simplified set of transdiagnostic symptom measures such that the same relationships can be tested in replication samples. Finally, we cannot exclude the possibility that treatment within the clinical cohorts may impact our results. Future studies examining drug-naïve populations, or with sample sizes sufficiently large to attempt to control for treatment type, will be important moving forwards.

## 6. Conclusion

The amalgamation of novel functional-molecular neuroimaging techniques, normative modelling, and a transdiagnostic perspective utilised here offers methodological and theoretical progress towards an understanding of the shared neurobiological foundations that underpin psychiatric conditions. Our transdiagnostic approach moves away from case-control analyses and offers an interesting way to situate clinical groups or individuals within between-subject similarity and deviation-symptom landscapes, which when scaled up across diagnoses, symptomatology, and molecular systems may offer novel perspectives on how complex aberrations of affect and cognition map onto dysfunction spanning molecular and systems level readouts. The long-term goal of this approach would be to build analytic bridges between these neuroimaging-derived brain phenotypes and treatments. Regardless of the progress made in diagnostics and prognostics, these both serve the ultimate goal of facilitating the provision of the right treatment to the right patient at the right time. The opportunity to link symptoms to non-invasive measures of molecular systems amenable to pharmacotherapeutic intervention may prove a useful tool for precision psychiatry in the longer term, helping to overcome the longstanding challenges of clinical heterogeneity and the hierarchical organisation of the brain.

## Acknowledgements of funding

TL is in receipt of a PhD studentship funded by the National Institute for Health Research (NIHR) Biomedical Research Centre at South London and Maudsley NHS Foundation Trust and King’s College London. DM, MAH, FET, and OD are supported by the NIHR Biomedical Research Centre and Clinical Research Facility at South London and Maudsley NHS Foundation Trust and King’s College London. MAH is also supported by the Medical Research Council (MR/N026969/1). MV is supported by EU funding within the MUR PNRR “National Center for HPC, BIG DATA AND QUANTUM COMPUTING (Project no. CN00000013 CN1), by the PNR National Grant DIGITAL LIFELONG PREVENTION (Project no PNC0000002_DARE), and by Wellcome Trust Digital Award (no. 215747/Z/19/Z). A.G. is supported by the KCL-funded CDT in Data-Driven Health; this represents independent research partly funded by the NIHR Maudsley’s Biomedical Research Centre (BRC) at the South London and Maudsley NHS Foundation Trust and partly funded by GSK. The views expressed are those of the authors and not necessarily those of the NHS, the NIHR, or the Department of Health and Social Care.

## Supplementary figures

**SI figure 1:**
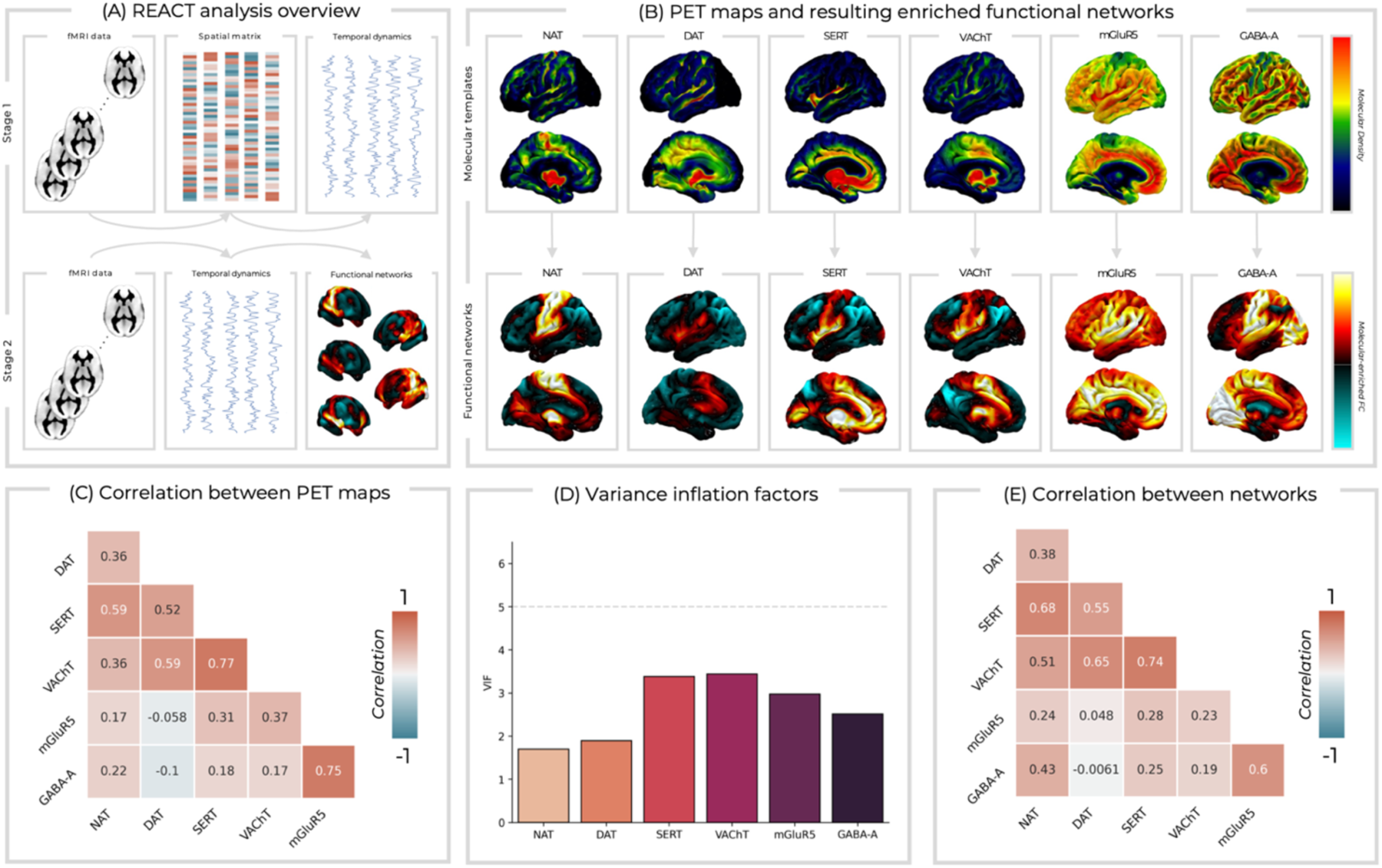
(A) An overview of the REACT methodology. (B) The PET maps utilised within the REACT analyses (top row) and the resultant molecular-enriched networks (bottom row). (C) Correlation coefficients between each pair of PET maps. (D) Variance inflation factors (VIF) for each PET map are below the rule of thumb value of 5, suggesting a non-problematic level of collinearity within the REACT model. (E) The correlation coefficients between each pair of molecular-enriched networks.

**Supplementary figure 2:**
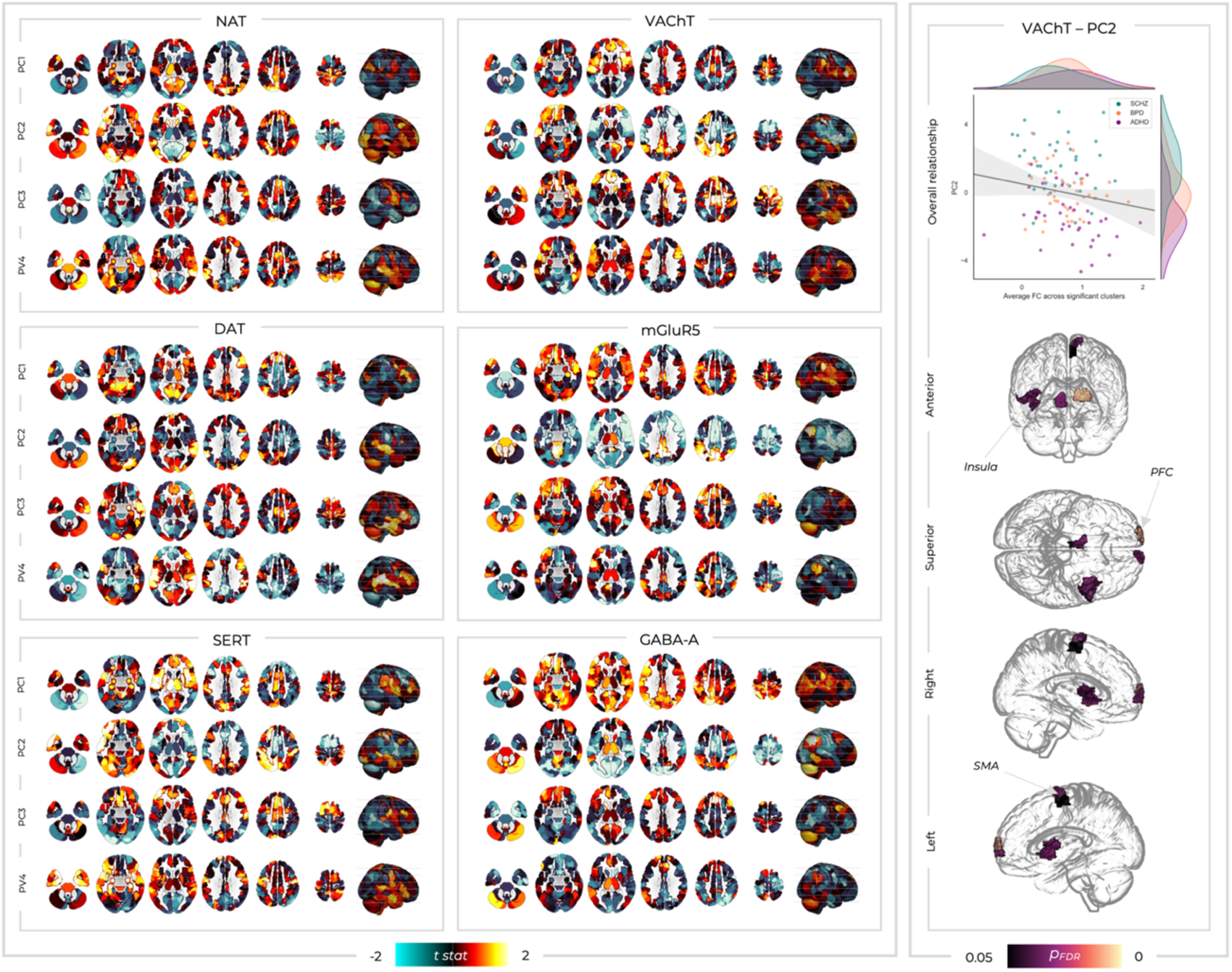
(Left box) The t statistics from multiple-linear regression analyses examining the relationship between molecular-enriched FC (as within a conventional REACT analysis) and the four principal components at each ROI. This allows for direct comparison with the normative modelling results running the same analysis but with deviations scores (figure X within main manuscript). (Right box) Only one of these deviation-symptom relationships was significant. Significant *p*_FDR_ values are shown for the relationships between PC2 and VAChT-enriched FC. The scatterplot shows this relationship with molecular-enriched FC averaged across all significant clusters. PFC; prefrontal cortex, SMA; supplementary motor area.

**S1 figure 3:**
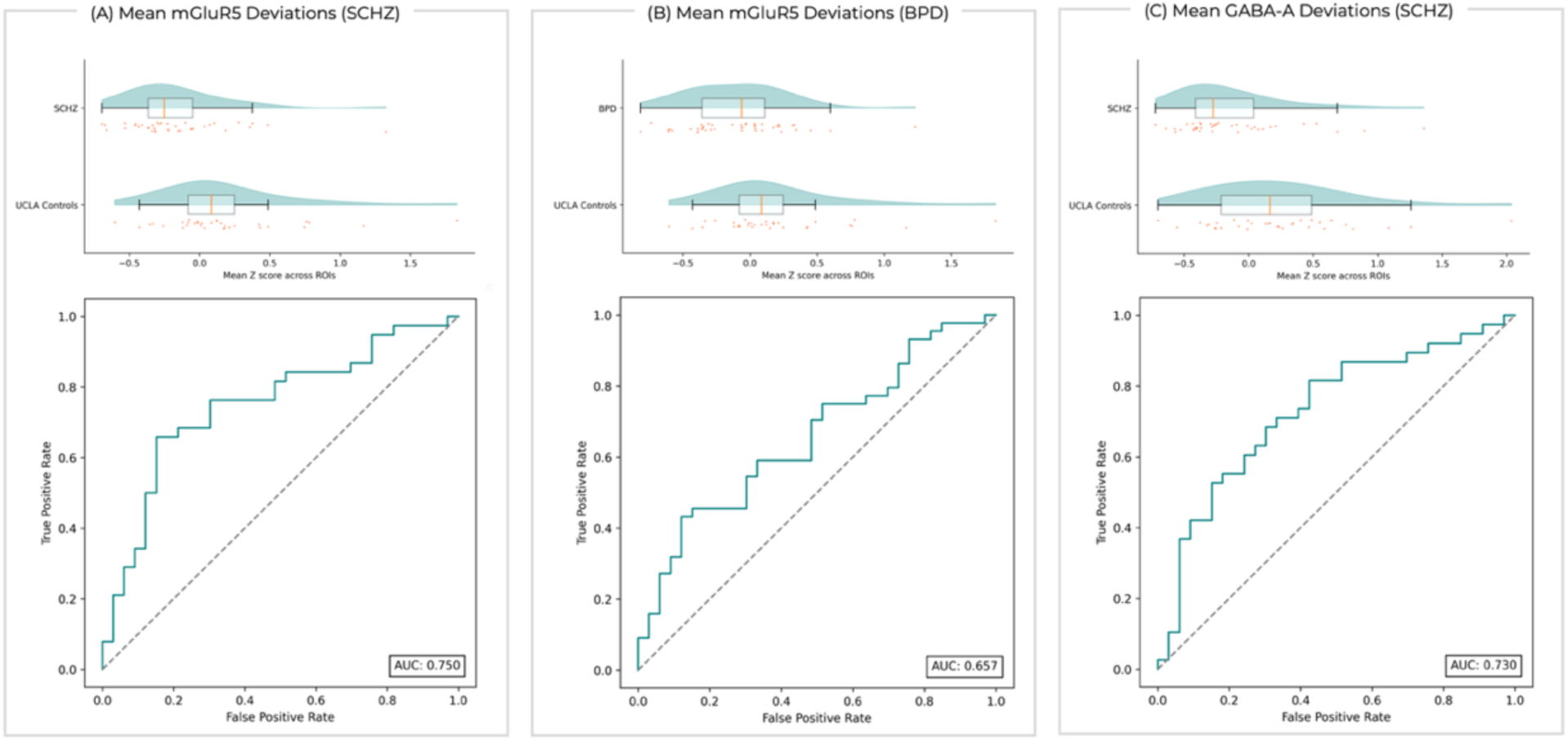
When comparing mean *Z* scores for each subject across the different groups, significant differences were found for mGluR5 between healthy individuals from the UCLA dataset and patients with SCHZ (A) and BPD (B). Similarly, healthy individuals from the UCLA dataset differed from SCHZ patients for the GABA-A system (C). The top row shows raincloud plots of these mean *Z* scores and the bottom row shows receiver operating curves demonstrating the predictive utility of these mean *Z* values for classifying patients from controls.

